# Primate homologs of mouse cortico-striatal circuits

**DOI:** 10.1101/834481

**Authors:** Joshua H. Balsters, Valerio Zerbi, Jérôme Sallet, Nicole Wenderoth, Rogier B. Mars

**Author notes:** Authors contributed equally to this project (Joshua H. Balsters, Valerio Zerbi). **Corresponding Author:** Dr Joshua Henk Balsters, Department of Psychology, Royal Holloway University of London, Egham, Surrey, UK, TW20 0EX.

## Abstract

With the increasing necessity of animal models in biomedical research, there is a vital need to harmonise findings across species by establishing similarities and differences in rodent and primate neuroanatomy. Using a connectivity fingerprint matching approach, we compared cortico-striatal circuits across humans, non-human primates, and mice using resting state fMRI data in all species. Our results suggest that the connectivity patterns for both the nucleus accumbens and cortico-striatal motor circuits (posterior/lateral putamen) were conserved across species, making them reliable targets for cross-species comparisons. However, a large number of human and macaque striatal voxels were not matched to any mouse cortico-striatal circuit (mouse->human: 85% unassigned; mouse->macaque 69% unassigned; macaque->human; 31% unassigned). These unassigned voxels were largely localised to the caudate nucleus and anterior putamen, overlapping with executive function and social/language regions of the striatum, and connected to prefrontal-projecting cerebellar lobules and anterior prefrontal cortex, forming circuits that seem to be unique for non-human primates and humans. Our results demonstrate the potential of connectivity fingerprint matching to bridge the gap between rodent and primate neuroanatomy.

## Introduction

Animal models are currently providing crucial insights into neural structure, function, and disorders. As the volume of non-human neuroscientific research increases, there is a growing necessity for translational, comparative neuroscience to harmonise these results with our understanding of structure, function, and disease in the human brain. To date, formal comparisons of brain organization across species have largely focussed on humans and non-human primates, in spite of the steady increase in rodent models in neuroscience (Carlén, 2017; Ellenbroek and Youn, 2016). This is partly driven by the lack of research in the different species using the same methods, but also by a distinct lack of consensus in terminology between research in rodents and research in primates, which is prohibitive of clear translation of results (Laubach et al., 2018). Here, we address the growing need to formally identify common brain circuits between rodents, non-human primates, and humans using the same technique to determine the scope and limits of rodent translational models.

One key issue that has hampered the comparison of rodent and primate brain organization has been establishing suitable comparative measures (Preuss, 1995). For instance, the discussion of the putative existence of a rodent homolog of human prefrontal cortex, different authors have proposed and dismissed a single connection to mediodorsal thalamus (Preuss, 1995; Rose and Woolsey, 1948), the presence of a granular layer IV (Preuss, 1995; Uylings et al., 2003), or equivalence of function (Dalley et al., 2004; Laubach et al., 2018) as diagnostics. These issues highlight the difficulties in understanding homologies across such distantly related species. Importantly, even if homology of areas is established, the homologous region will be embedded into a different large-scale network in the two species, which has consequences for the interpretation of translational results. Moreover, establishing similarity between anatomy and function is only useful if these two levels of description can be related. Ideally, we would want to understand what exactly is similar and different in the anatomical organization of the different species’ brains and how this relates to their behavioural abilities.

One approach to comparative neuroscience that has been successful at establishing similarities and differences on a continuous scale is that of matching areas across species based on their so-called ‘connectivity fingerprint’. The term connectivity fingerprint was introduced by Passingham et al (Passingham et al., 2002) to suggest that brain regions could be partitioned into functionally distinct brain areas based upon their unique set of connections, which in turn constrain the function of an area. For example, Passingham et al (Passingham et al., 2002) showed that macaque premotor areas F3 and F5 possess unique connectivity fingerprints, which correspond to their unique functional responses based on electrophysiology (neural firing during memory guided vs visually guided tasks respectively). Whilst connectivity fingerprints were previously used as a way to distinguish between functionally distinct regions within one brain, Mars et al (Mars et al., 2016) proposed that connectivity fingerprint matching could be used as a tool to identify similar brain areas across species. As case in point, Neubert et al (Neubert et al., 2014) systematically compared connectivity fingerprints from multiple regions in the human ventrolateral frontal cortex (vlFC) with connectivity fingerprints of regions defined in macaques. The results identified eleven vlFC brain regions with similar fingerprints in both species, and one brain region (lateral Frontal Pole; FPl) that was uniquely human. This approach has also been used to compare brain regions in the dorsolateral prefrontal cortex (PFC) (Sallet et al., 2013), medial PFC and orbitofrontal cortex (Neubert et al., 2015), parietal lobe (Mars et al., 2011), and temporoparietal cortex (Mars et al., 2013) in humans and non-human primates. However, it has to our knowledge never been used to compare neuroanatomical connectivity patterns across humans, non-human primates, and rodents.

This approach is now feasible due to the availability of the same type of neuroimaging data from humans, macaques, and rodents. Although tracer-based connectivity mapping is often considered to be the “gold standard” for comparative neuroscience, these methods are too expensive and labour intensive to use in most species and for the entire brain (Mars et al., 2014). The substantial investment of both time and money often means that it is only possible to investigate a small number of subjects which reduces the power of statistical comparisons and the ability to make global inferences. Also, these invasive approaches are far less common in humans, making it difficult to harmonise findings across species using a common methodology. Resting state fMRI (rsfMRI) is increasingly employed as a non-invasive tool to measure connectivity in humans and non-human primates (Neubert et al., 2015; Schaeffer et al., 2018; Vincent et al., 2007) and the availability of rodent resting state data (Grandjean et al., 2019; Zerbi et al., 2015) invites an extension of this work to larger-scale between-species comparisons. The increasing application of rsfMRI across species is likely because of the high degree of consistency between connectivity profiles derived from tracers and connectivity derived using rsfMRI (Grandjean et al., 2017). In addition, rsfMRI repositories are making large numbers of datasets freely available making it possible to conduct analyses using a common methodology across species with samples sizes that are orders of magnitude larger than traditional tracer-based approaches (Milham et al., 2018).

Here, we will use connectivity fingerprint matching to compare cortico-striatal circuits in humans, non-human primates, and rodents. The general architecture of cortical-striatal circuits, with partially separated loops connecting distinct parts of the striatum with distinct parts of the neocortex, seems well preserved across mammals (Haber, 2016; Heilbronner et al., 2016). However, the specific implementation can be expected to differ when the cortex has expanded in particular lineages, including the human (Murray et al., 2016). These circuits are particularly affected by a number of psychiatric conditions (Bradshaw and Enticott, 2001), some of which have been the target of recent rodent models (Zerbi et al., 2018). As such, cortico-striatal circuits are an ideal target to assess the feasibility of rodent-macaque-human translational studies.

## 2. Results

### 2.1. Establishing cortico-striatal connectivity fingerprints in mice

The connectivity fingerprinting approach requires extracting a rsfMRI timeseries from a seed region and comparing that with rsfMRI timeseries from a collection of target regions. The strength of the correlation between seed and target timeseries will be an indication of the strength of connectivity between those two regions, and the variability in connectivity across targets will produce the connectivity fingerprint. Mouse striatal seed regions were created using an independent tracer-based connectivity dataset from the Allen institute (Oh et al., 2014). The strength of tracer connectivity was established for each voxel in the mouse striatum and a hierarchical clustering algorithm was used to cluster voxels with similar and distinct connectivity patterns – an approach commonly referred to as connectivity-based parcellation (Balsters et al., 2016; 2018; Eickhoff et al., 2015). This approach identified three clusters in the mouse striatum with unique connectivity patterns based on tracer data; Nucleus Accumbens (NAcc), medial caudoputamen (mCP), and lateral caudoputamen (lCP) – figure 1a.

**Figure 1:**
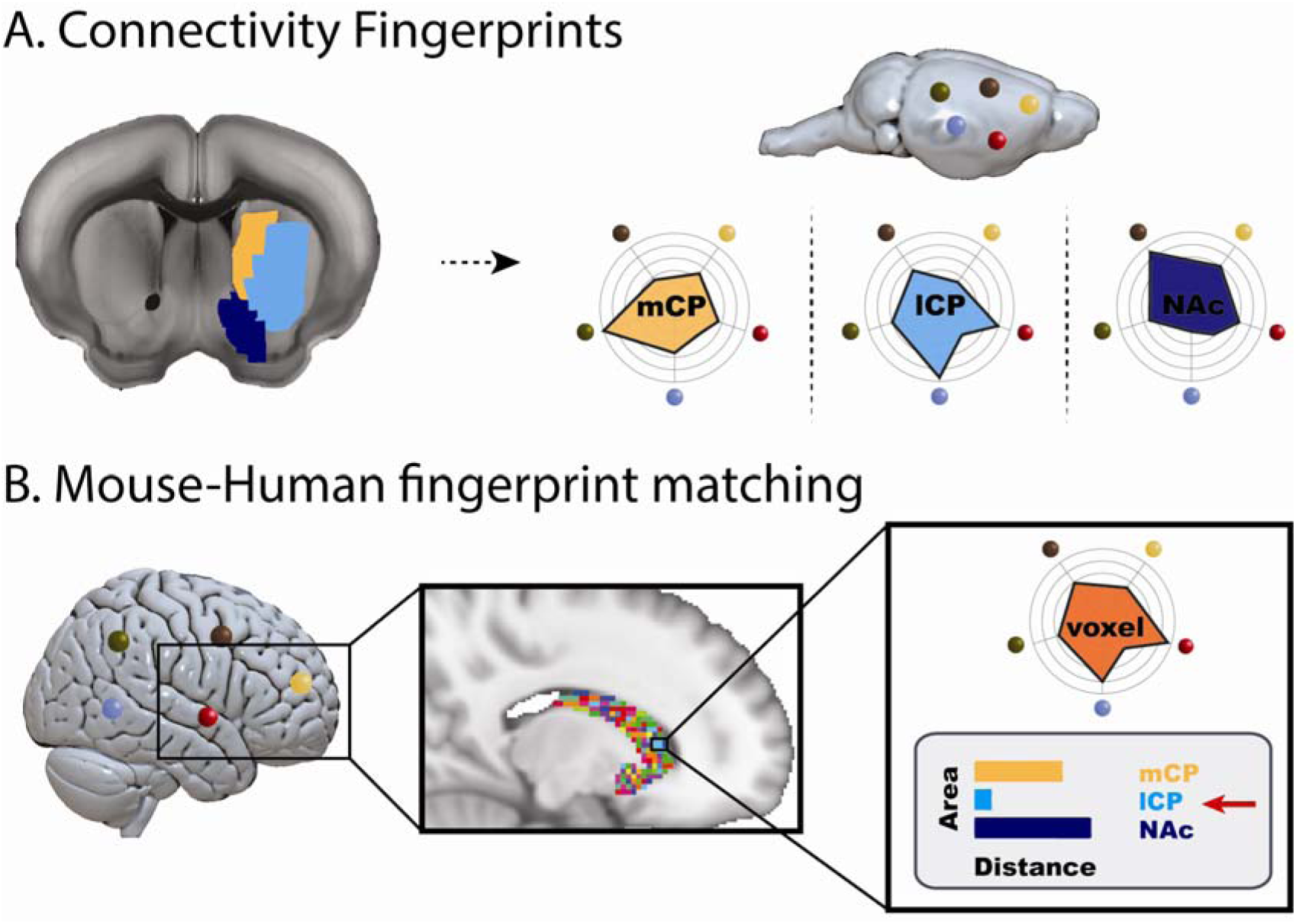
A schematic representation of the connectivity fingerprint matching procedure. A) The image left of the arrow shows the three cluster parcellation of the mouse striatum defined using tracer connectivity data from the Allen Brain Institute. The three clusters include the nucleus accumbens (NAcc; blue), medial caudoputamen (mCP: yellow), and lateral caudoputamen (lCP: cyan). Images right of the arrow show a schematic representation of how connectivity fingerprints were created for each of the three striatal seed regions. Connectivity fingerprints show the strength of connectivity (correlation of resting state fMRI timeseries) between each striatal seed region and target regions outside the striatum. B) In humans (or macaques), connectivity fingerprints were extracted from each voxel of the striatum - comparing the connectivity strength of striatal voxels with human homologs of the five target regions identified in mice. The similarity of each human voxel fingerprint can then be compared against each of the three mouse striatal fingerprints.

After creating mouse striatal seeds, we selected twelve target regions across cortical and subcortical regions of the mouse brain. These regions were chosen as they are believed to be homologous across species based on existing literature (see methods). We next compared connectivity fingerprints from the three striatal seed regions in an independent mouse rsfMRI dataset, extracting rsfMRI timeseries from the three striatal seeds and correlating these with timeseries extracted from the twelve target regions (figure 2). Analysis of the connectivity fingerprints (permutation testing of Manhattan distance – see methods) demonstrated that these three fingerprints were all significantly different from one another (mCP vs NAcc: Distance=2.57, p<0.001; mCP vs lCP: Distance=6.67, p<0.001; NAcc vs lCP: Distance=6.15, p<0.001). This confirms that we can extract unique connectivity patterns in the mouse striatum and thus have isolated suitable seeds and targets for testing similarities and differences in connectivity fingerprints across species. Voxel-wise correlation maps for these three seeds can also be seen in supplemental materials.

**Figure 2:**
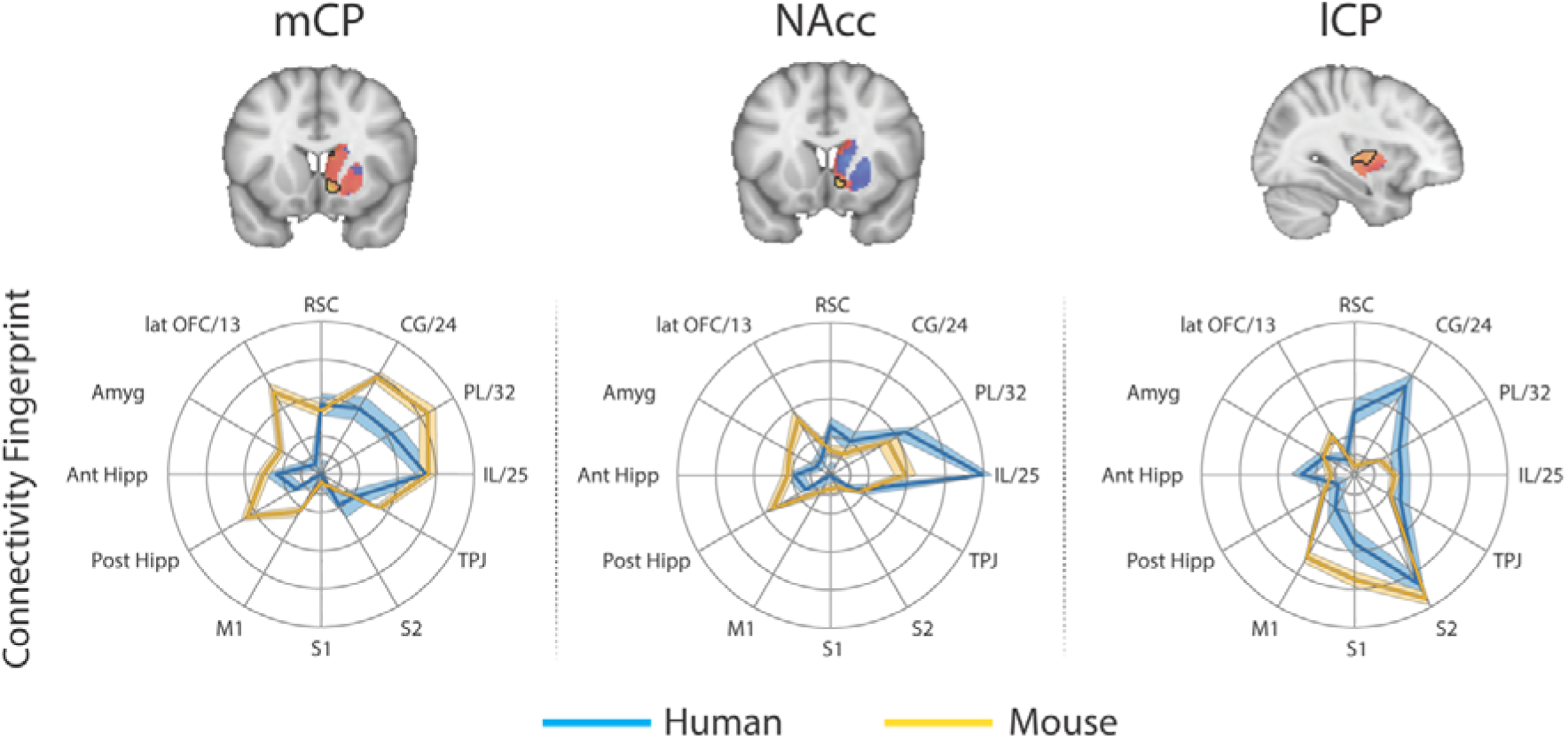
Brain images show unthresholded striatal t-maps of mouse-human similarity for mCP, NAcc, and lCP. Red-yellow voxels indicate increasingly positive correlations of connectivity fingerprints across species, blue-cyan voxels indicate increasing negative correlations of connectivity fingerprints across species. Black outlines indicate voxels that showed statistically significant similarity (TFCE p<0.05). Human and mouse connectivity fingerprints are shown underneath each brain image to highlight similarity in the connectivity pattern across species. Shaded error bars show the standard error of the mean. The data ranges from Z-values −0.1-0.4 and the thick grey circle shows 0.

### 2.2. Mouse to human comparison

These three mouse striatal seeds are commonly considered to reflect associative (mCP), limbic (NAcc), and motor circuits (lCP) (Hunnicutt et al., 2016). As such, we predicted that corresponding regions in the human striatum will exist within the human caudate nucleus, NAcc, and posterior putamen respectively. To test this, we extracted connectivity fingerprints for each voxel in the human striatum (defined as Harvard-Oxford subcortical atlas >33% threshold Caudate Nucleus, NAcc, and Putamen) and statistically compared each voxel’s connectivity fingerprint with the three mouse connectivity fingerprints (figure 1). This approach produced three voxelwise maps (Fishers r-to-z transformed maps) for each participant (one for each mouse fingerprint) illustrating voxelwise similarity with each mouse cortico-striatal circuit. These maps were input into a GLM (one sample t-test) for permutation testing and correction for multiple comparisons (TFCE p<0.05). All the unthresholded statistical maps in the section can be viewed at https://neurovault.org/collections/NFGTNVFX/. Voxels within the human NAcc possessed a statistically similar connectivity fingerprint with both the mouse NAcc and mCP, whereas connectivity fingerprints within human posterior putamen voxels were statistically similar to the mouse lCP (see figure 2). Figure 2 also highlights that even though there was an overlap in statistically similar voxels for mouse NAcc and mCP in the human NAcc, there were clear subthreshold differences in the human caudate and putamen (positive correlations with mCP but not NAcc) that distinguish these two striatal seeds. Using the task-based parcellation of Pauli et al (Pauli et al., 2016), we were able to assess the functional roles of the assigned human voxels. This showed that the human striatal voxels assigned to mCP and NAcc were best localised to regions of the striatum processing stimulus value, whereas the human striatal voxels assigned to lCP were localised to striatal regions contributing to motor control (see supplemental figure 3).

Whilst this approach identified common striatal regions in humans and mice, there were a high number of voxels in the human striatum (85%) that did not show a significant correspondence to any of the three striatal connectivity fingerprints from the mouse – we refer to these as unassigned voxels. Unassigned voxels accounted for 85% of the caudate nucleus volume, 77% of the putamen volume, and 5% of the NAcc volume. Functionally, the unassigned voxels were localised to striatal regions associated with executive function, social/language, and action value (Pauli et al., 2016). These unassigned voxels could reflect the expansion of the prefrontal cortex in primates.

To further establish the cortical-striatal connectivity of the human regions identified above, we next used the resulting similarity t-maps to extract weighted timeseries and correlated them with the connectivity pattern of each voxel in the rest of the brain (figure 3). Human mCP voxels showed significant connectivity with frontal pole regions (Area 47m), medial (RCZa, Area 23) and lateral prefrontal cortex (Area 46), anterior insula, supramarginal gyrus (hIP2), occipital pole (V1), cerebellar lobules HVI/Crus I and VIIIb, and hippocampus. The human homologue of NAcc also showed significant connectivity with anterior and middle cingulate cortex (Area 32dl and area 32p), occipital pole (V1), and cerebellar lobules Crus I and IX. The human homologue of lCP showed significant connectivity with the middle frontal gyrus (Area 9/46), precentral gyrus (areas 6 and 4p), SMA, Rolandic operculum (OP2), supramarginal gyrus (PFm), superior parietal lobule (area 7), precuneus (area 5), fusiform gyrus (FG2), and cerebellar lobule HVI. Full results tables are included in supplemental materials.

**Figure 3:**
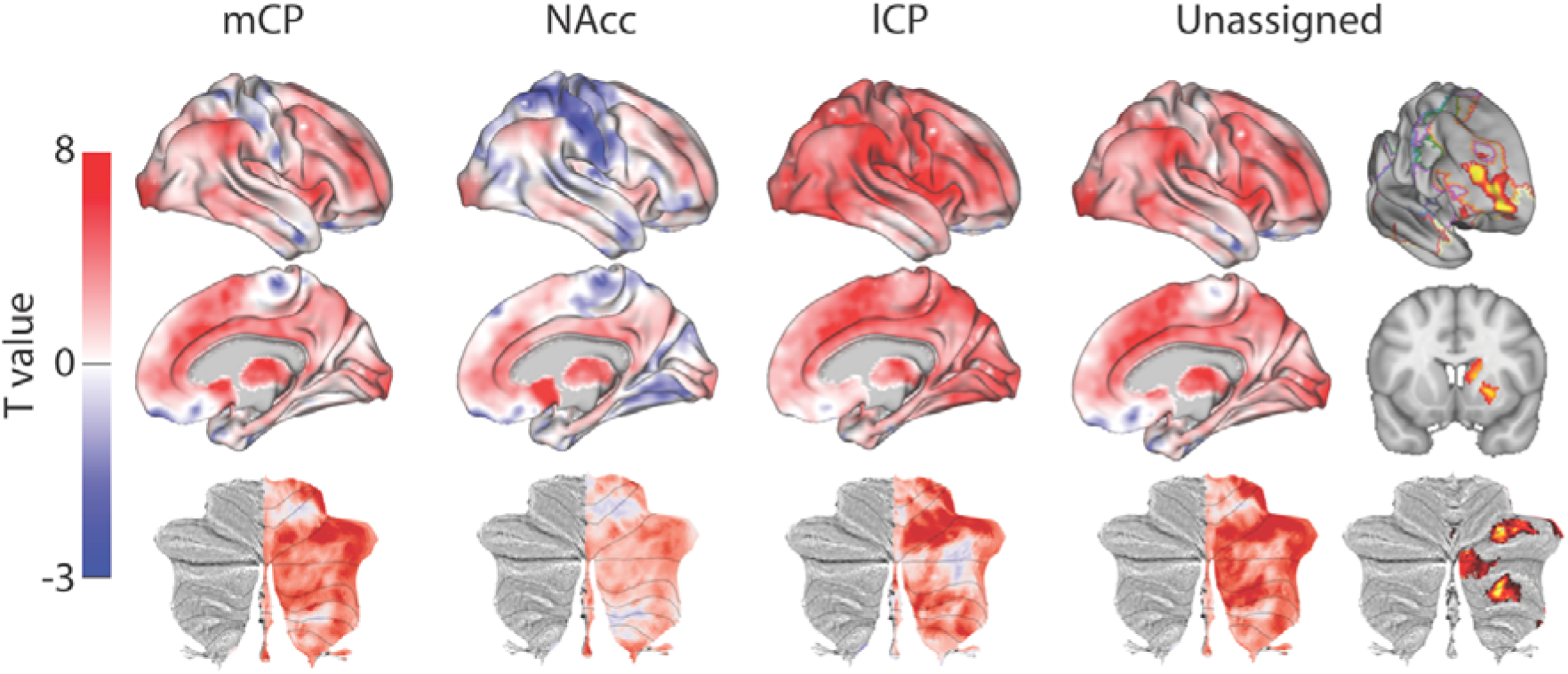
Unthresholded whole-brain connectivity maps showing regions interconnected with human homologs of mCP, NAcc, lCP, and unassigned voxels. The bottom row shows cerebellar activations on a flattened representation of the cerebellum and dotted black lines show the lobular boundaries (Diedrichsen and Zotow, 2015). The far-right column shows a thresholded conjunction analysis of voxels that possess significantly greater connectivity with unassigned voxels compared against all three mouse seeds. Outlines from the Yeo cortical parcellation highlight that the significantly different voxels in humans are principally in regions identified as the frontal -parietal network and the cerebellum.

In order to highlight significant differences between humans and mice (rather than relying the absence of a significant effect), we performed a conjunction analysis between unassigned whole-brain connectivity maps and whole-brain connectivity maps from the three mouse seeds (see figure 3). This analysis identified voxels in the human brain that showed significantly greater connectivity with unassigned voxels compared to all three human-mouse seeds independently (i.e. unassigned > mCP & unassigned > NAcc & unassigned > lCP). This analysis revealed striatal connectivity with the frontal pole (FPl and Area 46) was unique to humans. It also revealed structures known for their connections with the prefrontal cortex: mediodorsal nucleus of the thalamus, and cerebellar lobules Crus I and Crus II. Finally, this analysis also refined the localization of unassigned voxels within the human striatum. Whilst 85% of the human striatal voxels were not statistically similar to any of the three mouse cortico-striatal fingerprints, only 25.67% of those voxels were significantly different. These were localised to two distinct striatal subregions; the first in the rostral caudate nucleus, the second in the anterior portion of the putamen. Both of these striatal regions are associated with executive functions and language processes. Figure 3 shows that these unique cortico-striatal connections in humans also fall principally within the boundaries of the fronto-parietal network (FPN) as defined by Yeo et al (Yeo et al., 2011).

### 2.3. Mouse to macaque comparison

We next applied the same procedure to compare cortico-striatal connectivity fingerprints in mice with cortico-striatal connectivity fingerprints in the macaque. Both species underwent similar rs-fMRI protocols including light sedation using anaesthesia, and as such this comparison allows us to account for the potential effects of anaesthesia on rs-fMRI connectivity. We hypothesised that a similar number of voxels will be unassigned in this comparison given that, from an evolutionary perspective rodents and primates diverged around 89M years ago.

For each voxel in the macaque striatum (defined as caudate nucleus, NAcc, and putamen maps from the INIA19 macaque atlas; Rohlfing et al (Rohlfing et al., 2012)) we extracted a connectivity fingerprint using the same twelve targets. Figure 4 shows that once again, we found regions of significant similarity, specifically the mCP and NAcc showed significant similarity with voxels in the macaque NAcc, caudate head and caudate tail. The lCP showed significant overlap with posterior regions of the putamen. This left 69% of voxels in the macaque striatum unassigned, i.e. that did not show significant similarity with any mouse connectivity fingerprints. Unassigned voxels accounted for 79% of the caudate nucleus volume, 62% of the putamen volume, and 9% of the NAcc volume.

**Figure 4:**
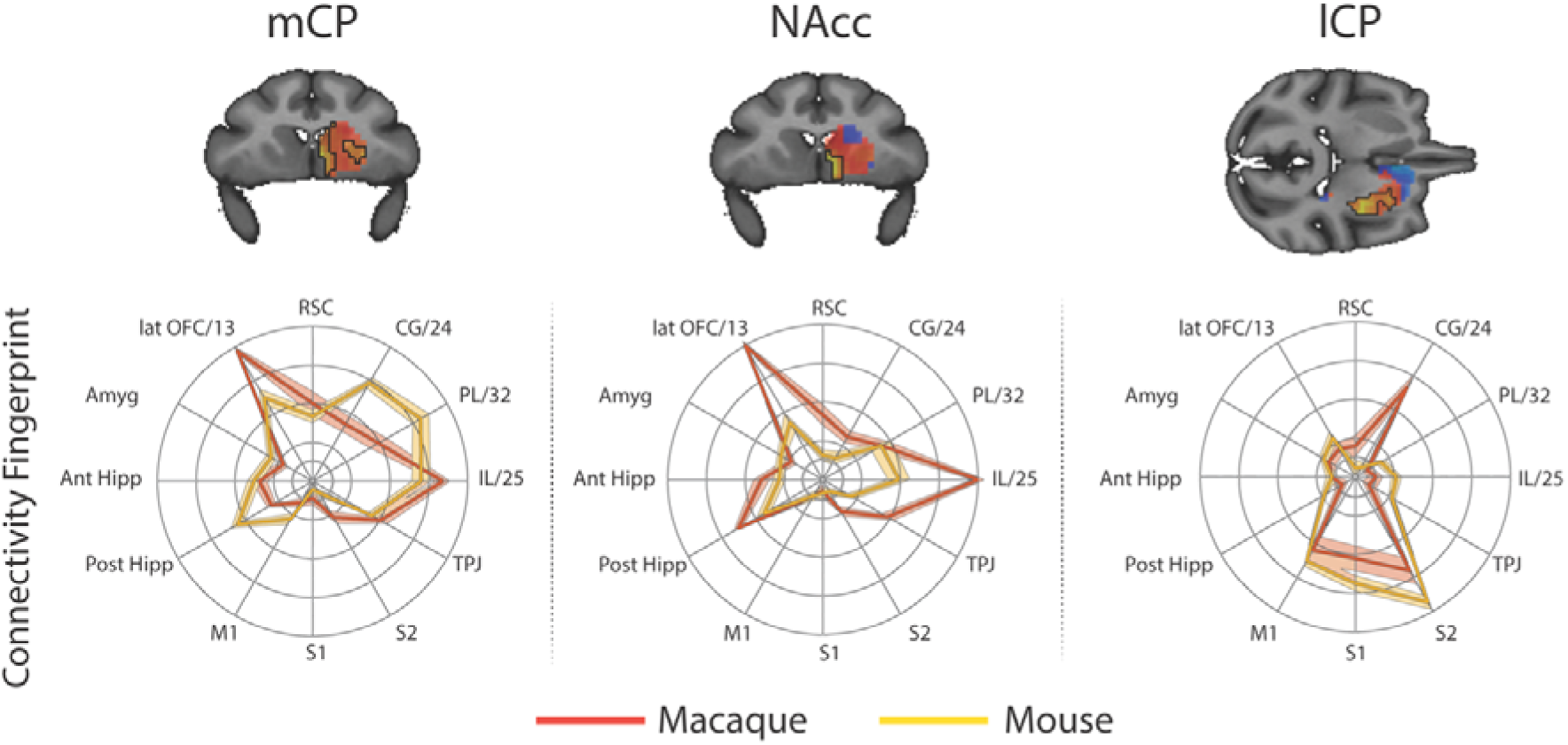
Brain images show unthresholded striatal t-maps of mouse-macaque similarity for mCP, NAcc, and lCP. Red-yellow voxels indicate increasingly positive correlations of connectivity fingerprints across species, blue-cyan voxels indicate increasing negative correlations of connectivity fingerprints across species. Black outlines indicate voxels that showed statistically significant similarity (TFCE p<0.05). Macaque and mouse connectivity fingerprints are shown underneath each brain image to highlight the similarity in the connectivity pattern across species. Shaded error bars show the standard error of the mean. The data ranges from Z-values −0.1-0.4 and the thick grey circle shows 0.

Figure 5 displays regions showing significant connectivity with mCP, NAcc, and lCP voxels. MCP voxels showed significant connectivity with the dorsomedial prefrontal cortex (Areas 9m), premotor cortex (F2, F5), posterior lateral prefrontal cortex (Area 45b), anterior cingulate cortex (Area 24), posterior cingulate cortex (Area 23b), intraparietal sulcus (LIPd), cortex of the superior temporal sulcus and visual areas (V1 and V2). Significant connectivity with subcortical structures included the amygdala and caudate nucleus. Regions showing significant connectivity with NAcc voxels included anterior cingulate cortex (Area 24), premotor cortex (F4 and F5), posterior lateral prefrontal cortex (area 44), cortex of the superior temporal sulcus, and visual cortex (V2). Subcortical connectivity included the macaque NAcc, amygdala, and hippocampus. Regions showing significant connectivity with lCP voxels included the premotor cortex (F2, F5), anterior cingulate cortex (area 24) and posterior cingulate cortex (Areas 23 and 31), somatosensory cortex on the posterior bank of central sulcus, and Intraparietal sulcus (LIP). Subcortical regions included the putamen (largest activation) and the Caudate nucleus.

**Figure 5:**
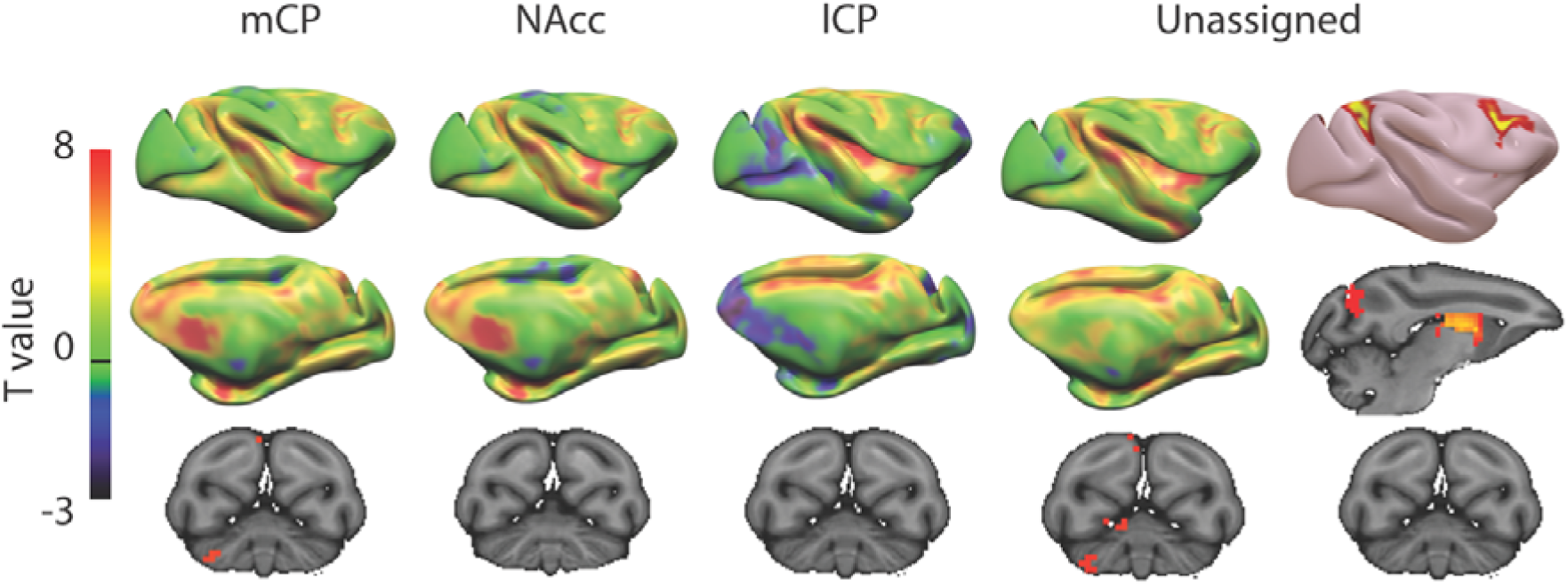
Unthresholded whole-brain connectivity maps showing regions interconnected with mCP, NAcc, lCP, and unassigned voxels. The bottom row shows cerebellar activations. The far-right column shows a thresholded conjunction analysis of voxels that possess significantly greater connectivity with unassigned voxels compared against all three mouse seeds.

The conjunction analysis (i.e. unassigned > mCP & unassigned > NAcc & unassigned > lCP) highlighted regions of significantly different in macaques compared to mice. These included dorsolateral prefrontal cortex (Area 46d), premotor cortex (F2), Anterior Insula, cortex of the superior temporal sulcus, inferior parietal lobule (Area 7A), and visual cortex (V1). Subcortical differences were seen in both the rostral caudate nucleus and putamen as in humans. The application of the conjunction analysis showed that only 20% of striatal unassigned voxels were significantly different from all three mouse seeds. These significantly different voxels accounted for 34% of the caudate nucleus, 10% of the putamen, and 1% of NAcc volume (4 voxels), suggesting that differences between macaques and mice were largely driven by differences in the caudate nucleus - as in the previous mouse-human analyses. Full results tables are included in supplemental materials.

### 2.4. Macaque to Human comparisons

In order to determine whether some striatal connectivity features of the unassigned voxels could be reflecting uniquely human specializations, we applied the same protocols to compare striatal connectivity fingerprints in humans and macaques. Similar to what was done for the mouse, macaque caudate body, NAcc, and putamen were defined as seeds using connectivity-based parcellation (see methods). We used two target models; 1) the original targets used in the mouse model and 2) an extended model that included additional regions in the lateral and medial PFC that have been shown previously to have homologous connectivity fingerprints across macaques and humans (Neubert et al., 2014; Sallet et al., 2013). This included area 9/46d, area 9/46v, Area 44, area FPm, and SMA. Connectivity fingerprints for all three seeds were independent in both the mouse model (Caudate vs NAcc: Distance=3.51, p<0.001; Caudate vs Putamen: Distance=3.62, p<0.001; NAcc vs Putamen: Distance=4.96, p<0.001) and the extended model (Caudate vs NAcc: Distance=4.77, p<0.001; Caudate vs Putamen: Distance=5.32, p<0.001; NAcc vs Putamen: Distance=6.12, p<0.001).

For each voxel of the human Caudate Nucleus, NAcc, and Putamen (based on Harvard Oxford subcortical atlas >33% threshold) we extracted the human connectivity fingerprint and compared it to connectivity fingerprints for the macaque caudate body, NAcc, and putamen. Unsurprisingly, the human-macaque comparisons were more closely aligned then the mouse-human comparisons. Specifically, the human voxels in the caudate nucleus were significantly similar to the macaque caudate body, human voxels in the NAcc were significantly similar to the macaque NAcc, and human voxels in the putamen were significantly similar to the macaque putamen (see figure 6). Using the task-based parcellation of Pauli et al (Pauli et al., 2016), the human voxels assigned to the macaque caudate body overlapped with striatal regions dedicated to executive function, social/language processes, and action value. The human voxels assigned to the macaque NAcc overlapped with striatal regions dedicated to stimulus value processing. The human voxels assigned to the macaque putamen best overlapped with striatal motor control regions (see supplemental figure 6).

**Figure 6:**
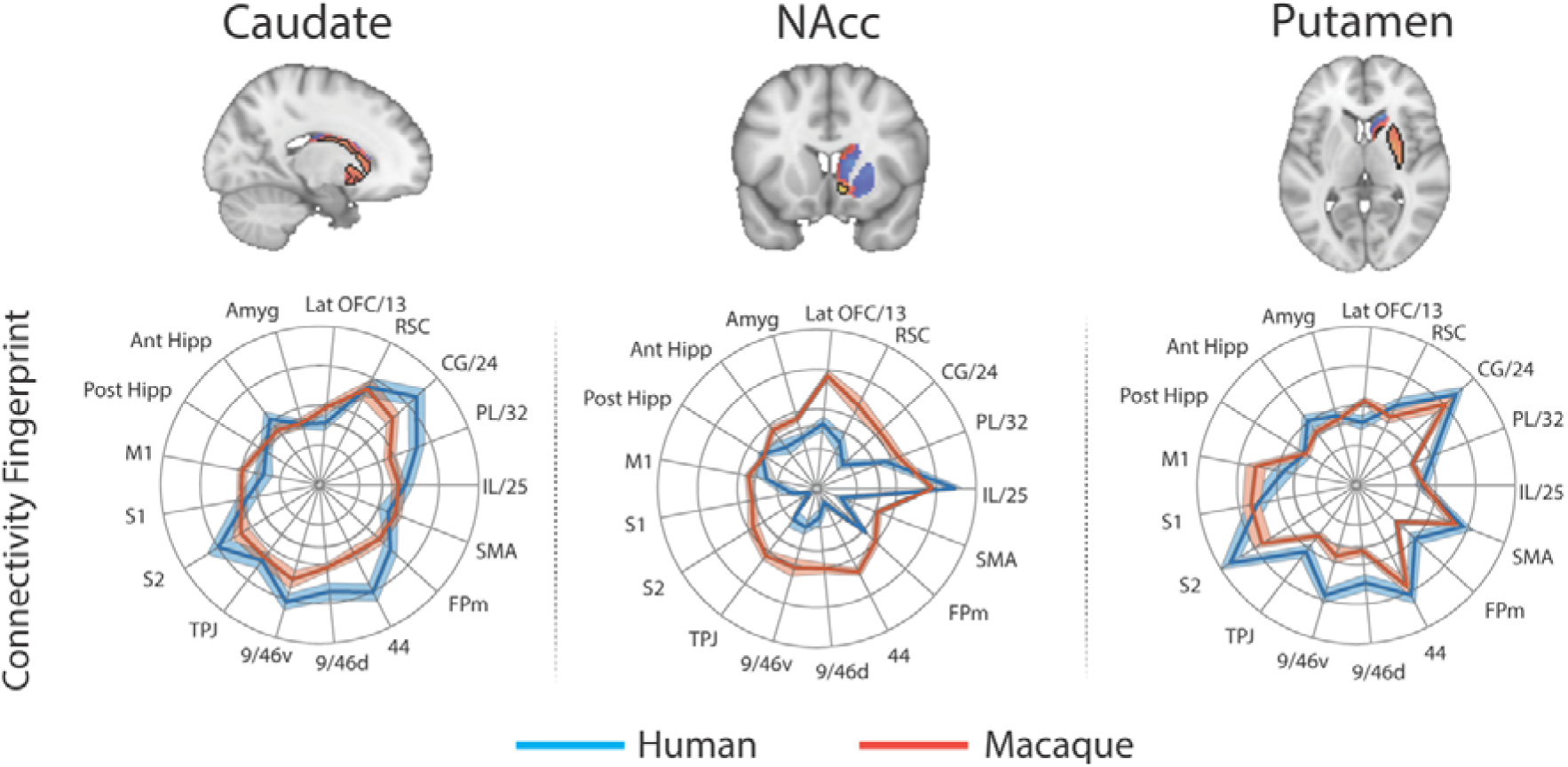
Brain images show unthresholded striatal t-maps of macaque-human similarity for caudate body, NAcc, and putamen. Red-yellow voxels indicate increasingly positive correlations of connectivity fingerprints across species, blue-cyan voxels indicate increasing negative correlations of connectivity fingerprints across species. Black outlines indicate voxels that showed statistically significant similarity (TFCE p<0.05). Human and macaque connectivity fingerprints are shown underneath each brain image to highlight the similarity in connectivity pattern across species. Shaded error bars show the standard error of the mean. The data ranges from Z-values −0.25-0.4 and the thick grey circle shows 0.

The human-macaque comparison produced fewer unassigned voxels than the human-mouse comparison, with only 31% of striatal voxels unassigned using the same model used in mice, and 20% unassigned using the extended model. These voxels were localised anatomically to the NAcc (30% of NAcc voxels unassigned using the reduced mouse model, 24% using the extended model) and Caudate (35% of caudate voxels unassigned using the reduced mouse model, 23% using the extended model), and showed functional association with striatal regions associated with executive functions. Although 30/23% of unassigned voxels were localised in the NAcc, the conjunction analysis isolating significant differences between species showed that only voxels in the caudate nucleus were significantly different between species.

As previously, we extracted weighted timeseries from each similarity t-map to investigate voxelwise connectivity (figure 7). The human-macaque caudate nucleus maps showed significant connectivity with mid cingulate cortex (RCZa, CCZ) and middle (Area 46) and inferior frontal gyri (pars triangularis), inferior parietal lobule (hIP3), precuneus (Area 7p), occipital pole (V1 and V3), cerebellar lobules HVI and Crus II. Human-macaque NAcc maps showed significant connectivity with the anterior cingulate cortex (area 32pl) and cerebellar lobule Crus IX. Human-macaque putamen maps showed significant connectivity with regions in the mid cingulate cortex (RCZa) and lateral prefrontal cortex (area 46) and premotor cortex (area 6), supramarginal gyrus (area PF), middle temporal gyrus, occipital lobe (V3), and cerebellar lobule HVI.

**Figure 7:**
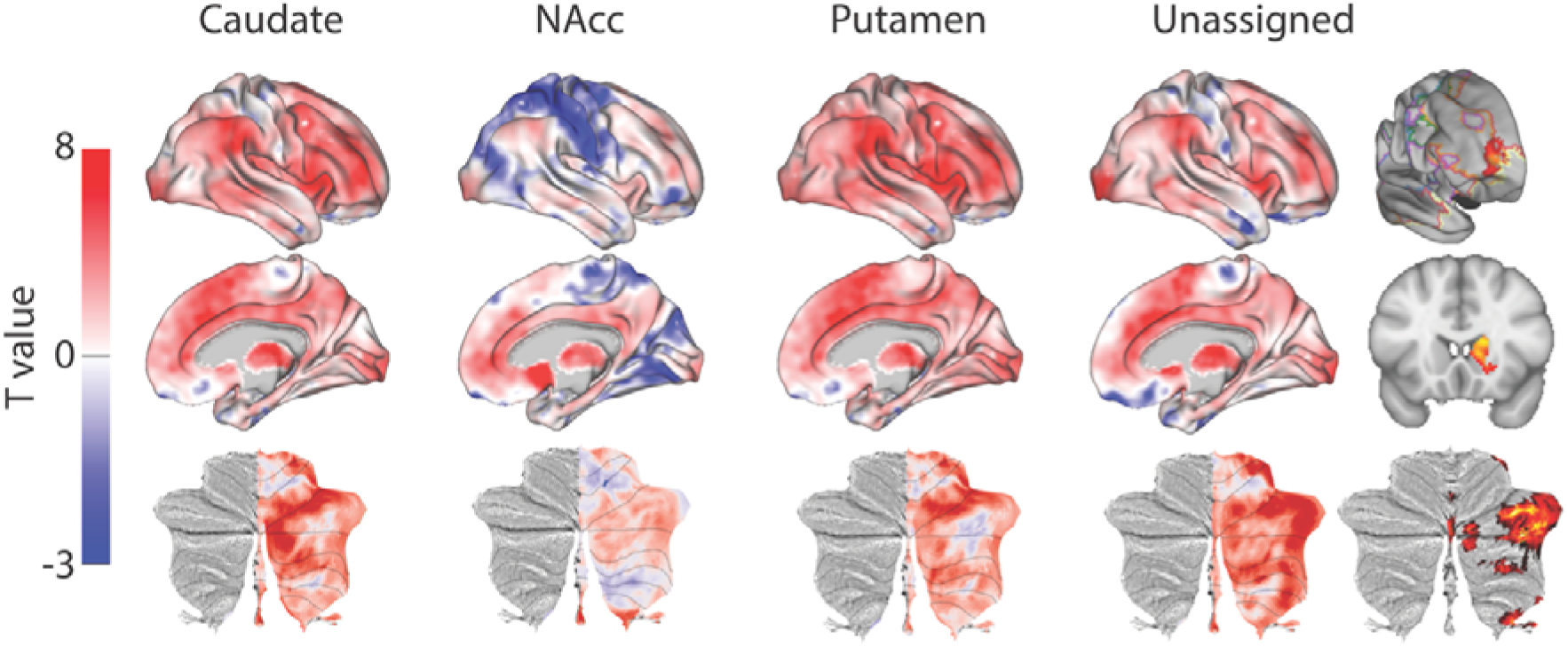
Unthresholded whole-brain connectivity maps showing regions interconnected with caudate body, NAcc, putamen, and unassigned voxels. The bottom row shows cerebellar activations on a flattened representation of the cerebellum and dotted black lines show the lobular boundaries (Diedrichsen and Zotow, 2015). The far-right column shows a thresholded conjunction analysis of voxels that possess significantly greater connectivity with unassigned voxels compared against all three mouse seeds. Outlines from the Yeo cortical parcellation highlight that the significantly different voxels in humans are principally in regions identified as the frontal -parietal network and the cerebellum.

When comparing connectivity differences between assigned vs unassigned voxels using the conjunction analysis, the unassigned voxels showed significant connectivity with frontal pole (FPl), precuneus, occipital pole (V1), and subcortical structures known for projecting to the prefrontal cortex (the mediodorsal nucleus of the thalamus and the cerebellar lobule Crus I). Coupling with the FPl could reflect the expansion of the lateral frontal pole since the last common ancestor to humans and macaques (Neubert et al. 2014). The number of unassigned voxels was similar to the number of voxels that were significantly different (31% unassigned voxels but 18.1% of them were significantly different to our 3 fingerprints). Although unassigned voxels were localised to both the NAcc and caudate nucleus, only voxels in the caudate nucleus were significantly different in humans compared to macaques. These have been localised to regions of caudate connected with the FPN that process executive functions and action value. As with mice, this unique cluster of voxels was co-localised with FPN as defined by Yeo et al (Yeo et al., 2011). All the statistical maps reported in the section can be viewed at https://neurovault.org/collections/NFGTNVFX/.

## 3. Discussion

As the use of mouse models for studying function and disease rapidly increases in neuroscience, it is crucial to develop methods that can harmonise results across species. Here for the first time we have used rsfMRI as a common methodology to establish similarities and differences in striatal-cortical organization in humans, non-human primates, and rodents. Using our connectivity fingerprint matching approach, we could identify NAcc consistently across species making it a reliable target for translational neuroscience. Although portions of the caudate nucleus and putamen in both humans and macaques showed similar connectivity fingerprints with the mouse, there were also large regions of the human and macaque striatum that appeared to be unique and unassigned. Regions of significant difference across species appear to be mostly localised to the anterior putamen and caudate body. In both human-mouse and human-macaque comparisons, unassigned voxels showed significantly greater connectivity with the lateral frontal pole (Area 46 in mice and FPl in both mice and macaques) and prefrontal-projecting subcortical structures including the dorsomedial nucleus of the thalamus and cerebellar lobules Crus I and Crus II.

### Striatum

Consistent with previous parcellations of the mouse striatum, our connectivity based parcellation using Allen tracer data identified three regions with unique connectivity fingerprints; the NAcc, medial and lateral CP. Previous studies have suggested that these regions map on to the distinct functional domains of limbic, association, and sensorimotor respectively (Balleine et al., 2009; Hunnicutt et al., 2016; Thorn et al., 2010; Yin and Knowlton, 2006). Our results suggest that the connectivity fingerprint of the NAcc is highly conserved across rodents and primates. Human and macaque striatal voxels assigned to the mouse NAcc, and the human voxels assigned to the macaque NAcc, were all discretely localised within the boundaries of the human NAcc as defined by the Harvard-Oxford subcortical atlas and INIA atlas in primates (see figure 8). This is consistent with tracer studies by Mailly et al (Mailly et al., 2013) and Heilbronner et al (Heilbronner et al., 2016) who also demonstrated an overlap in NAcc connectivity fingerprints across rodents and non-human primates. This conserved brain network could underpin their similar functional role in motivation and reinforcement learning. For example, lesions to the rat NAcc have been shown to alter performance in value-based decision-making paradigms such as delayed discounting tasks (Cardinal et al., 2001). Similarly, non-invasive imaging studies in humans have linked individual variability in NAcc activity to individual variability in delayed discounting preferences, mirroring what has been shown in rodents (Hariri et al., 2006). Aberrant performance on value-based decision-making paradigms, as well as altered NAcc activity, has been proposed as a hallmark for a number of psychiatric conditions including Obsessive Compulsive Disorder, addiction, schizophrenia and others (Gunaydin and Kreitzer, 2015). Given the high degree of conservation across species in NAcc connectivity, we would suggest that the NAcc is a reliable translational target for rodent models of psychiatric conditions.

**Figure 8:**
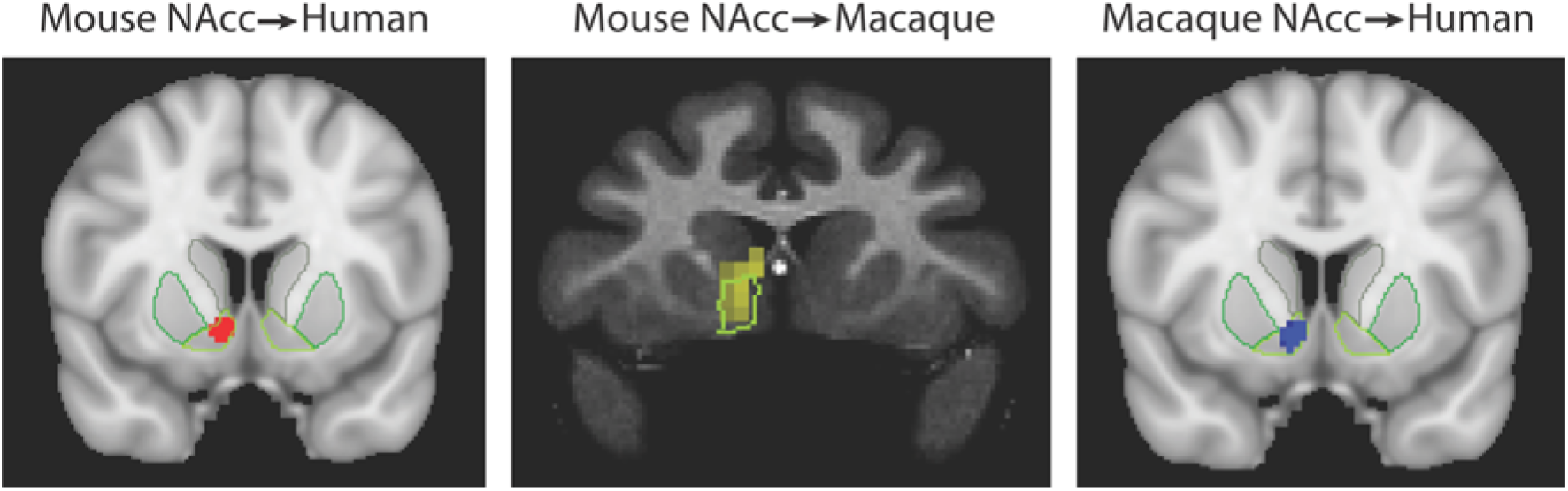
Brain images showing the spatial overlap of the NAcc across species. Voxels highlight regions of statistically significant similarity across species (TFCE p<0.05). The light green outline in each image shows the anatomical boundaries of the human and macaque NAcc.

The mCP and lCP are believed to contribute to associative and sensorimotor motor processes respectively and would thus be expected to correspond to regions of the caudate and putamen with similar functional properties. Our results showed that voxels in the posterior segment of the putamen in both humans and macaques shared a significantly similar connectivity fingerprint with the mouse lCP (see figures 2 and 4), suggesting that sensorimotor cortico-striatal circuits could also be comparable across species. This is crucial for translational models of sensorimotor deficits such as Parkinson’s disease. However, 85% of voxels in the human striatum failed to show significant similarity with any of the three mouse striatal seeds. These unassigned voxels were localised to the caudate nucleus and putamen. The conjunction analysis highlighted voxels that were significantly different in humans compared to mice (as opposed to voxels that failed to reach the statistical significance threshold). This approach confirmed that 25% of voxels in human striatum – specifically the anterior portion of the putamen and caudate body - possess significantly different connectivity fingerprints in humans compared to mice. These regions of the striatum have been shown to receive projections from the lateral prefrontal cortex in both humans and non-human primates (Alexander et al., 1986; Choi et al., 2017; Verstynen et al., 2012), and our comparison with task-based parcellations of the striatum suggest that these regions process action value, executive functions, and social and language processes in a meta-analysis of over 10,000 human fMRI studies. Although these results could reflect the effects of anaesthesia on rs-fMRI in mice, we believe this is unlikely given that 1) significant similarities were found between humans and mice for limbic (NAcc) and sensorimotor (lCP) cortico-striatal networks, and 2) rs-fMRI data collected from macaques used a similar anaesthesia protocol to that of mice and yet the macaque caudate nucleus showed significant similarity with most voxels in the human caudate nucleus (figure 6). We suggest that these differences between human and mouse striatal fingerprints are more likely to reflect differences in connectivity with the lateral prefrontal cortex, and as such we would caution researchers using rodent models for disorders affecting predominantly executive function and social/language functions as there appears to be less clear translation between rodents and humans.

One caveat to the approach used here is the definition of striatal seeds in the mouse. Although three subdivisions of the rodent striatum are largely consistent with previous studies (Balleine et al., 2009; Hunnicutt et al., 2016; Thorn et al., 2010; Yin and Knowlton, 2006), it is possible that an increasingly fine grained parcellation would produce more precise human homologs. Hintiryan et al (Hintiryan et al., 2016) utilised a novel neuroanatomical approach to map the cortico-striatal projectome in mice, identifying 29 striatal regions with distinct connectivity fingerprints. The limitations of fMRI and its spatial resolution make such a detailed analysis unfeasible, however, with increasing advances in MRI methods it may be possible in the future to investigate higher resolution parcellations of the striatum and other cortical and subcortical structures. We note, however, that the current data are of the best resolution and quality currently in general use (Grandjean et al., 2017; Marcus et al., 2016; Milham et al., 2018) and it is likely most translational studies will not be working with data of superior quality.

### Prefrontal cortex

Most theories on the functions of the prefrontal cortex have been derived from studies in humans and nonhuman primates, awaiting translation in rodents. Multiple approaches have been taken to establish homologs across species, including similarities in cytoarchitecture, similarities in connectivity, and equivalence of function. While each of these approaches has been used to debate the existence of the granular PFC in rodents, there is at least increasing consensus that rodent prefrontal regions IL, PL, and Cg could be equivalent to human and macaque areas 25, 32, and 24 respectively (Bicks et al., 2015). Heilbronner et al (Heilbronner et al., 2016) showed similar striatal projection patterns in rodents and macaques using tracers from IL/25, PL/32, and Cg/24 in both these species. However, there still is quite some debate on whether other parts of human prefrontal cortex are present in the rodent (Laubach et al., 2018). At the very least, human prefrontal cortex as extended substantially in absolute terms, although its relative extension compared to other primates is also a matter of fierce debate (Barton and Venditti, 2013; Passingham and Smaers, 2014). For the definition of our connectivity fingerprints, we have only used regions whose homology across species has been well established in the literature (see methods and supplemental tables 1-3). However, our results speak to the debate about translational results in prefrontal cortex in two ways.

First, and most obvious, we find voxels in the human striatum that possess a connectivity fingerprint that is not found in either the mouse or macaque. These voxels all tended to have a strong functional connectivity with, among others, parts of the human lateral prefrontal cortex (area 46 and lateral frontal pole (FPl)). Although area 46 has a clear homolog in the macaque, it is part of granular dorsolateral prefrontal cortex which is believed to be an anthropoid primate specialization (Passingham and Wise, 2012; Preuss and Goldman-Rakic, 1991) and therefore not present in the mouse. The lateral part of the frontal pole has been identified in humans based on both cytoarchitecture and connectivity (Bludau et al., 2014), but a macaque homolog is not detectable using rs-fMRI raising the possibility that this is a human or at least ape specialization (Neubert et al., 2014). The fact that the unassigned striatal voxels predominantly show connectivity with these areas suggests the presence of a unique cortical-striatal loop. Based on our findings, this loop would be mostly involved in higher-order human behaviours, as are associated with FPl (Hartogsveld et al., 2018; Vendetti and Bunge, 2014), suggesting that these might be difficult to study using the translational paradigm.

Our findings also warrant a second caution with regard to the translation of prefrontal results. Even though human striatal regions possessed similar connectivity fingerprints to those of mice, it is possible that similar circuits could include novel projections in humans (Mars et al., 2018). For instance, voxels in the human striatum assigned to mCP showed the expected connectivity with homologous target regions such as the medial frontal cortex, but also with regions in the dorsal prefrontal cortex (dlPFC) and inferior parietal lobule (Figure 3). Since it’s disputed whether homologs of dlPFC and inferior parietal lobule exist within mice, it seems likely the mCP network is embedded within a larger network in humans that includes novel regions. We therefore urge caution interpreting translational results: even if regions of interest are homologous, they might be embedded in larger networks that include novel areas.

### Cerebellum

Humans showed unique striatal connectivity with parts of association cortex, including the anterior prefrontal cortex, but also with large regions of cerebellar cortex. Specifically, the cerebellar lobules showing unique striatal connectivity in humans were Crus I and Crus II, which have been shown to be interconnected with the prefrontal cortex (Kelly and Strick, 2003; O’Reilly et al., 2010), and contribute to cognitive processes including rule-guided behaviour (Balsters and Ramnani, 2008; 2011; Balsters et al., 2013) and language (Lesage et al., 2012; 2017; Mariën et al., 2014). Tracer studies in rodents and non-human primates have shown that prefrontal projections to the cerebellar cortex are conserved across species (Kelly and Strick, 2003; Schmahmann and Pandya, 1997; Wiesendanger and Wiesendanger, 1982), however research also suggests that prefrontal-cerebellar circuits have selectively expanded in humans compared to other species. Specifically, prefrontal projections to the pontine nucleus (Ramnani et al., 2006), volume of prefrontal-projecting cerebellar lobules Crus I and Crus II (Balsters et al., 2010; Luo et al., 2017), and the volume and connections of the dorsal dentate nucleus (Baizer, 2014; Matano, 2001; Steele et al., 2016) have all expanded in humans relative to motor cerebellar circuitry across species. Studies of cerebellar evolution have generally suggested that Crus I and Crus II are homologous across species (Larsell, 1952) even though they have expanded in humans. The findings of this study could suggest that along with selective expansion, there may be additionally novel cortico-cerebellar connections in humans that have contributed to the expansion of the cortico-cerebellar system in humans. Given that the Area FPl homolog in non-human primates and rodents is unclear or absent, it is plausible that this region has generated novel projections to both the striatum and cerebellum. The finer grained 17-Network parcellation of the human cerebral cortex by Yeo et al (Yeo et al., 2011), includes a region similar to Area FPl identified in this study (Network 13). Buckner et al’s (Buckner et al., 2011) analysis of cerebellar connectivity using the Yeo cortical parcellation shows that this region projects to Crus I and Crus II, adding further evidence to suggest that the some of the expansion of Crus I and Crus II could be due to novel inputs from the frontal pole in humans.

### Conclusions and cautions

Here, we demonstrate the potential of connectivity fingerprint matching to bridge the gap between rodent and primate neuroanatomy. Our results highlight the core properties of a rodent to primate striatum, including similarities in connectivity fingerprints for NAcc across species that could be a useful model for translational neuroscience. However, we would caution against researchers comparing medial and lateral regions of the mouse caudoputamen with the primate caudate nucleus and putamen. Although homologs were identified, there were also clear differences in connectivity patterns that require further investigation. We propose that these differences reflect the expansion of frontal cortex in primates, along with the relative expansion of area FPl in humans. These results will hopefully add to the on-going debates surrounding similarities in cortical brain regions across species, i.e. the existence of the PFC in rodents. Further studies using connectivity fingerprint matching could help to refine where similarities and differences exist across species in other brain structures including medial prefrontal cortex and orbitofrontal regions.

## 4. Methods

### 4.1. Targets & seeds

#### 4.1.1. Reduced/Mouse model

The reduced model includes twelve target regions common to all species: 1) Infralimbic (Area 25), 2) Prelimbic (Area 32), 3) Cingulate areas (Area 24), 4) Retrosplenial Cortex (Area 30), 5) Lateral Orbitofrontal Cortex (Area 13), 6) Basolateral Amgydala, 7) Dorsal (Anterior) Hippocampus, 8) Ventral (Posterior) Hippocampus, 9) Primary Motor Cortex (M1), 10) Primary Sensory Cortex (S1), 11) Supplemental Sensory Cortex (S2), 12) Temporal association area (TPJp). Targets were 3×3×3 voxels in all species. Further details can be found in supplemental tables 1-3.

#### 4.1.2. Extended/Primate model

The extended model includes the twelve reduced model targets and five additional targets that are common to macaques and humans, but not mice: 13) Area 9/46d, 14) Area 9/46v, 15) Area 44d, 16) FPm, 17) SMA. Targets were 3×3×3 voxels in all species. Further details can be found in supplemental tables 2 and 3.

#### 4.1.3. Mouse striatal seeds

Anterograde viral-tracer maps were obtained using the query form from the Allen Institute database and resampled at 100 µm^3^ to match the fMRI data resolution. Individual experiments were selected as follows: carried in wild-type C57BL/6 and with injection volume >0.1µl. The connectivity was determined from the injection site to the projections by quantifying the fluorescence locally for each voxel (Grandjean et al., 2017; Oh et al., 2014).

We established connectivity strength (Z-transformed terminal tracer volume) between 68 injection sites in isocortex and each voxel in the caudoputamen, NAcc, and fundus. We then used connectivity-based parcellation to partition the mouse basal ganglia into regions with unique connectivity fingerprints (Balsters et al., 2016, 2018). The optimal solution based on silhouette value was three regions with unique connectivity fingerprints based on anterograde tracers. These are labelled medial caudoputamen (mCP), Nucleus accumbens (NAcc), and lateral caudoputamen (lCP). The percent variance explained by the first eigenvariate for each seed was 57.72% ± 6.42, 49.18% ± 6.12, and 60.14% ± 6.89 for mCP, NAcc, and lCP respectively.

#### 4.1.4. Macaque seeds

Connectivity-based parcellation was applied to the macaque resting state data in order to create seeds with unique connectivity fingerprints. The optimal solution based on silhouette value was a 5 cluster solution keeping the NAcc and putamen whole, and segmenting the caudate nucleus into 3 segments (the body and 2 segments in the tail of the caudate). We focussed our analysis on the caudate body, NAcc, and putamen given the small number of voxels contributing to the caudate tail seeds. The percent variance explained by the first eigenvariate for each seed was equivalent to that of mice - 56.14% ± 6.39, 51.83% ± 5.87, and 56.28% ± 7.56 for Caudate body, NAcc, and putamen respectively.

### 4.2. MRI data acquisition

#### 4.2.1. Mouse

Mouse fMRI and anatomical scans were collected from 20 wildtype C57BL/6J animals (males, median age = 82 days; median weight 26 grams). Animals were caged in standard housing, with food and water *ad libitum*, and a 12h day/night cycle. Protocols for animal care, magnetic resonance imaging, and anesthesia were carried out under the authority of personal and project licenses in accordance with the Swiss federal guidelines for the use of animals in research, and under licensing from the Zürich Cantonal veterinary office.

Anesthesia was induced with 4% isoflurane and the animals were endotracheally intubated and the tail vein cannulated. Mice were positioned on a MRI-compatible cradle, and artificially ventilated at 80 breaths per minute, 1:4 O_2_ to air ratio, and 1.8⍰ml/h flow (CWE, Ardmore, USA). A bolus injection of medetomidine 0.05⍰mg/kg and pancuronium bromide 0.2⍰mg/kg was administered, and isoflurane was reduced to 1%. After 5⍰min, an infusion of medetomidine 0.1⍰mg/kg/h and pancuronium bromide 0.4⍰mg/kg/h was administered, and isoflurane was further reduced to 0.5%. The animal temperature was monitored using a rectal thermometer probe, and maintained at 36.5⍰°C⍰±⍰0.5 during the measurements. The preparation of the animals did not exceed 20⍰min.

Data acquisition was performed on a Biospec 70/16 small animal MR system (Bruker BioSpin MRI, Ettlingen, Germany) with a cryogenic quadrature surface coil (Bruker BioSpin AG, Fällanden, Switzerland). After standard adjustments, shim gradients were optimized using mapshim protocol, with an ellipsoid reference volume covering the whole brain. For functional connectivity acquisition, a standard gradient-echo EPI sequence (GE-EPI, repetition time TR⍰=⍰1000⍰ms, echo time TE⍰=⍰15⍰ms, in-plane resolution RES⍰=⍰0.22⍰×⍰0.2⍰mm^2^, number of slice NS = 20, slice thickness ST = 0.4 mm, slice gap SG = 0.1 mm) was applied to acquire 2000 vol in 38 min. In addition, we acquired anatomical T2*-weighted images (FLASH sequence, in-plane resolution of 0.05 × 0.02 mm, TE = 3.51, TR = 522 ms). The levels of anesthesia and mouse physiological parameters were monitored following an established protocol to obtain a reliable measurement of functional connectivity (Grandjean et al., 2014; Zerbi et al., 2015).

#### 4.2.2. Macaque

Macaque fMRI and anatomical scans were collected from 10 healthy macaque monkeys (*Macaca mulatta*, 10 males, median age = 4.98 years; median weight 9.25 kg). Protocols for animal care, magnetic resonance imaging, and anaesthesia were carried out under the authority of personal and project licenses in accordance with the UK Animals (Scientific Procedures) Act 1986 (ASPA).

Anaesthesia was induced using intramuscular injection of ketamine (10 mg/kg) either combined with xylazine (0.125-0.25 mg/kg) or with midazolam (0.1 mg/kg) and buprenorphine (0.01mg/kg). Macaques also received injections of atropine (0.05 mg/kg intramuscularly), meloxicam (0.2 mg/kg intravenously) and ranitidine (0.05 mg/kg intravenously). Anaesthesia was maintained with isoflurane. The anesthetized animals were either placed in an MRI compatible stereotactic frame (Crist Instrument Co., Hagerstown, MA, USA) or resting on a custom-made mouth mold (Rogue Research, Mtl, QC, CA). All animals were then brought in a horizontal 3T MRI scanner with a full-size bore. Resting-state fMRI data collection commenced approximately 4 hours after anaesthesia induction, when the peak effect of ketamine was unlikely to be still present. In accordance with veterinary instruction, anaesthesia was maintained using the lowest possible concentration of isoflurane gas. The depth of anaesthesia was assessed using physiological parameters (continuous monitoring of heart rate and blood pressure as well as clinical checks for muscle relaxation prior to scanning). During the acquisition of the MRI data, the median expired isoflurane concentration was 1.083% (ranging between 0.6% and 1.317%). Isoflurane was selected for the scans as resting-state networks have previously been demonstrated to closely match known anatomical circuits using this agent (Neubert et al., 2014; Vincent et al., 2007). Slight individual differences in physiology cause slight differences in the amount of anaesthetic gas concentrations needed to impose a similar level of anaesthesia on different monkeys.

All but one animal were maintained with intermittent positive pressure ventilation to ensure a constant respiration rate during the functional scan; one macaque was breathing without assistance. Respiration rate, inspired and expired CO_2_, and inspired and expired isoflurane concentration were monitored and recorded using VitalMonitor software (Vetronic Services Ltd., Devon). In addition to these parameters, core temperature was monitored using a Opsens temperature sensor (Opsens, Quebec, Canada) and pulse rate, SpO_2_ (> 95%) were monitored using a Nonin sensor (Nonin Mediacal Inc., Minnesota, USA) throughout the scan.

A four-channel phased-array radio-frequency coil in conjunction with a local transmission coil was used for data acquisition (Dr. H. Kolster, Windmiller Kolster Scientific, Fresno, CA, USA). Whole-brain blood oxygen level dependent (BOLD) fMRI data were collected for 1600 volumes from each animal (except for one with 950 volumes), using the following parameters: 36 axial slices, in-plane resolution 1.5×1.5 mm, slice thickness 1.5 mm, no slice gap, TR=2280 ms, TE=30 ms. Structural scans with a 0.5mm isotropic resolution were acquired for each macaque in the same session, using a T1-weighted MP-RAGE sequence.

#### 4.2.3. Human

Twenty volumetric (as opposed to grayordinate) resting state fMRI datasets (Age 22-35yrs; 13 male) were downloaded from the Human Connectome Project (HCP) (Essen et al., 2013). Whole-brain BOLD EPI images were collected for approximately 15mins (1200 volumes) using a standardised protocol (2mm isotropic resolution, 72 slices, TR=720ms, TE=33.1ms, multiband factor=8). Only the first session with phase encoding left-to-right (LR) was used. Rs-fMRI data were already pre-processed using the FIX pipeline (automatic ICA rejection and regression of 24 head motion parameters) (Griffanti et al., 2014; Salimi-Khorshidi et al., 2014) and normalised into MNI space.

### 4.3. Resting state pre-processing and analysis

In all species, we used an ICA nuisance regression pre-processing strategy (FIX for human and mouse, manual component rejection for macaque). For all species, resting state analyses were conducted using CONN (Whitfield-Gabrieli and Nieto-Castanon, 2012). Data were bandpass filtered according to recent recommendations (Mouse: 0.01-0.25Hz; Macaque: 0.01-0.087Hz; Human: 0.01-0.15Hz), linear detrended, and despiked. Lowpass filter limits were set to be 5 volumes of data in all species, however this was further reduced for the human data in order to avoid physiological artefacts which are believed to occur at frequencies >0.2Hz (Baria et al., 2011).

Resting state analyses began with the creation of three template connectivity fingerprints in mice (mCP, NAcc, lCP) and macaques (caudate, NAcc, and putamen). For each dataset, we extracted the principle eigenvariate from three striatal seeds and correlated these with timeseries extracted from target regions. To confirm that of each striatal seed has a unique connectivity fingerprint we used the MrCat toolbox (https://github.com/neuroecology/MrCat) to establish the Manhattan distance between striatal connectivity fingerprints within species (i.e. comparing mCP and lCP fingerprints in mice) and used permutation testing (10,000 permutations) to test for significance. This same procedure was used to confirm that macaque striatal connectivity fingerprints were significantly different from one another.

Individual connectivity fingerprints within species were averaged (robust mean) to create template connectivity fingerprints for each striatal seed (see figures 2,4,6). In the comparison species, we extracted a connectivity fingerprint for each striatal voxel (correlation between the voxel and target timeseries). This voxel-based fingerprint was correlated with each of the template fingerprints from a different species and the resulting correlation value assigned to the voxel. This produced three correlation maps for each participant (one for each striatal seed) describing the correlation between each voxel fingerprint and the template fingerprint. Maps were Fisher’s r-to-Z transformed and run through permutation testing in FSL’s randomise (10,000 permutations, TFCE corrected p<0.05) to establish which voxels showed a significantly similar connectivity fingerprint.

T-Maps generated in the previous step were used as ROIs in CONN to generate whole-brain connectivity maps. A weighted timeseries from only positive (i.e. similar) voxels was extracted and used to identify connected regions across all grey matter voxels within the right hemisphere. Significant connectivity was established using permutation testing (10,000 permutations) and correction for multiple comparisons (p<0.001 voxel threshold, cluster-extent p<0.05 FDR). A conjunction analysis was employed to compare connectivity maps for assigned and unassigned voxels (Friston et al., 2005; Price and Friston, 1997). This approach is a more stringent comparison as it requires connectivity in the unassigned voxel map to be significantly greater than the connectivity for all of the assigned voxel maps (i.e. unassigned > mCP && unassigned > NAcc && unassigned > lCP).

### 4.4. Anatomical and functional localization

Mouse-to-human and macaque-to-human striatal homologs were localised using the Harvard Oxford subcortical atlas and a task-based parcellation of the striatum (Pauli et al., 2016). Rather than using the traditional approach of localisation based on local maxima, we used the distribution-based cluster assignment method outlined in Eickhoff et al (Eickhoff et al., 2007) to highlight the central tendency of activations and avoid localising to peripheral structures. This approach compares probability distributions for the underlying anatomical regions with probability distributions for the functional activation cluster, allowing one to make judgments about whether brain regions are over or under-represented. Specifically, the mean probability for area X at the location of the functional activation is divided by the overall mean probability for area X in all voxels where it was observed. This provides a quotient which indicates how much more (or less) likely an area was observed in the functionally defined volume than could be expected if the probabilities at that location would follow their overall distribution. A quotient > 1 indicates a rather central location of the activation with respect to this area, whereas a quotient < 1 a more peripheral one. Cortical activations were localised using a combination of cytoarchitectonic probability maps from the Anatomy Toolbox (Eickhoff et al., 2005; 2006; 2007) and connectivity-based parcellation maps available in FSLEYES (Mars et al., 2011; 2013; Neubert et al., 2014; 2015; Sallet et al., 2013; Tziortzi et al., 2014). Cerebellar activations were localised using the probabilistic cerebellar atlas (Diedrichsen et al., 2009).

## Acknowledgements

V.Z. is supported by an SNSF AMBIZIONE grant PZ00P3_173984/1. J.S is supported by a Wellcome Trust grant 105651/Z/14/Z. N.W. is supported by ETH Research Grant ETH-38 16-2. R.B.M. is supported by the Biotechnology and Biological Sciences Research Council (BBSRC) UK [BB/N019814/1] and the Netherlands Organization for Scientific Research NWO [452-13-015]. The Wellcome Centre for Integrative Neuroimaging is supported by core funding from the Wellcome Trust [203139/Z/16/Z].

